# On the Non-uniqueness Problem in Integrated Information Theory

**DOI:** 10.1101/2021.04.07.438793

**Authors:** Jake R. Hanson, Sara I. Walker

## Abstract

Integrated Information Theory is currently the leading mathematical theory of consciousness. The core of the theory relies on the calculation of a scalar mathematical measure of consciousness, Φ, which is deduced from the phenomenological axioms of the theory. Here, we show that despite its widespread use, Φ is not a well-defined mathematical concept in the sense that the value it specifies is neither unique nor specific. This problem, occasionally referred to as “undetermined qualia”, is the result of degeneracies in the optimization routine used to calculate Φ, which leads to ambiguities in determining the consciousness of systems under study. As demonstration, we first apply the mathematical definition of Φ to a simple `AND+OR` logic gate system and show 83 non-unique Φ values result, spanning a substantial portion of the range of possibilities. We then introduce a Python package called `PyPhi-Spectrum` which, unlike currently available packages, delivers the entire spectrum of possible Φ values for a given system. We apply this to a variety of examples of recently published calculations of Φ and show how virtually all Φ values from the sampled literature are chosen arbitrarily from a set of non-unique possibilities, the full range of which often includes both conscious and unconscious predictions. Lastly, we review proposed solutions to this degeneracy problem, and find none to provide a satisfactory solution, either because they fail to specify a unique Φ value or yield Φ = 0 for systems that are clearly integrated. We conclude with a discussion of requirements moving forward for scientifically valid theories of consciousness that avoid these degeneracy issues.

## 1 Introduction

Integrated Information Theory (IIT) is a leading contender among theories of consciousness due to its provision of a scalar mathematical measure, Φ, posited to predict the overall level of consciousness in virtually any dynamical system. In comparison to other contemporary theories of consciousness, such as Global Neuronal Workspace Theory [1] or Predictive Processing [2–6], IIT is set apart by its mathematical rigor. The concrete, mathematical formulation of the theory is at least partially responsible for its widespread adoption over the past decade^1^. However, there is some debate about IIT’s mathematical implementation. One of the more widely-discussed problems is the ambiguity in the derivation of Φ from the phenomenological axioms of the theory [8–10]. Not often discussed nor addressed is that the value of Φ one predicts is not unique, even within the standard accepted mathematical definition of Φ put forth in IIT 3.0 [11]. Indeed, as we will show, IIT can simultaneously predict a system to be both conscious and unconscious, implying the theory is ill-defined.

It has been recognized since as early as 2012 that Φ may be indeterminate for some systems [12]. Yet, to date, an investigation of how this impacts conclusions that can be drawn from the theory across different systems it is applied to has not been undertaken. Here we show, even the simplest of systems can evade a straightforward evaluation of the mathematical value of Φ. The specifics of this problem result as a consequence of what has been referred to as “underdetermined qualia” or “tied purviews” [13, 14]. In short, the problem results as a consequence of the fact that Φ (“big Phi”) is defined as an information-theoretic distance between cause-effect structures (vectors) that are not guaranteed to be unique. The non-uniqueness of cause-effect structures is, in turn, caused by an inability to select between degenerate core causes and effects (probability distributions) based on their *ϕ* (“little phi”) values. According to IIT 3.0, the cause/effect repertoire with the smallest *ϕ* value must be identified as the core cause/effect to satisfy the exclusion axiom but, in practice, there is no guarantee that this element is unique.

A standard resolution to this degeneracy problem is to select a core cause and core effect repertoire arbitrarily from the set of possibilities. For example, using an `if less than` statement in the optimization routine will select the first of the degenerate core causes/effects while using an `if less than or equal to` statement will select the last of the degenerate core causes/effects. In the configuration files for the software package `PyPhi` [15], which is used ubiquitously for the calculation of Φ [16–25], there is an option to select whether to keep the smallest or largest purview element in the event of a tie (i.e., the same *ϕ* value for different causes/effects). However, this choice is ad hoc for obvious reasons (see, *e.g.*, [13, 14, 26]): while it appears to provide a unique Φ value despite degeneracy, it is not a valid solution [14] because the tied purview elements are often the same size, meaning an element must still be selected arbitrarily. In this case, `PyPhi` defaults to selecting the first of the degenerate values which depends arbitrarily on the order in which elements are considered. A second significant issue with this solution is that there is nothing phenomenological built into the theory to suggest why the smallest or largest purview element should be retained as the unique cause/effect. Consequently, different authors have reached conflicting conclusions as to whether the smallest or largest purview element is more in line with the phenomenological axioms of the theory [13, 14, 26].

Here, we aim to shed light on these issues by calculating Φ for a very simple model system in the form of an `AND` gate connected to an `OR` gate. Taking the mathematical definition of Φ at face value, we demonstrate that a spectrum of 83 different Φ values results, corresponding to both conscious and unconscious predictions. Next, we provide a modified version of `PyPhi` called `PyPhi-Spectrum` that can be used to calculate the entire spectrum of Φ values for a given dynamical system with a single function call, as opposed to the singular value that is typically reported. We then apply this algorithm to a corpus of ten recently published Φ values, in order to determine the extent to which non-unique Φ values are overlooked in the literature. Last, we investigate whether or not proposed solutions adequately address this problem. We conclude with a philosophy of science discussion related to the scientific handling of ideas.

## 2 Methods

In what follows, we assume the reader is familiar with the basic terms in IIT 3.0. For a more detailed explanation of terms, we refer the reader to the original IIT 3.0 publication [11].

### 2.1 Preliminaries

In IIT 3.0, Φ^*Max*^ is the overall level of conscious experience that is predicted for a given dynamical system. The calculation of Φ^*Max*^ is notoriously difficult to perform. In total, five nested optimization steps are required, as shown in Algorithm 1. At the core of this routine is a simple distance measure in the form of an earth mover’s distance [27] between two probability distributions. This results in a measure of integration known as *ϕ* (“little phi”). However, this elementary distance calculation must be performed for every possible partition of a given “purview” in order to calculate *ϕ^MIP^*, then every possible purview for a given “mechanism” in order to find *ϕ^Max^*. This results in what is known as a cause-effect structure (CES) or “constellation”, which is defined by the set of mechanisms, their *ϕ^Max^* values, and two probability distributions per mechanism corresponding to the “core cause” and “core effect”. Next, one must generate a constellation for every possible partition of the subsystem under consideration and use a modification of the earth mover’s distance to quantify how close this constellation is to that of the unpartitioned subsystem. This results in a second measure of integration known as Φ (“big Phi”), which is designed to quantify the effect of a system-level partition on the underlying ability for a system’s components (mechanisms) to integrate information. The system-level partition with the smallest Φ value is the minimum information partition (MIP) and the corresponding Φ value is Φ^*MIP*^. Last, this entire process must be repeated for every possible subsystem in a given system in order to find the maximum integrated information Φ^*Max*^. In total, this hierarchy of nested optimization routines results in a computational complexity that scales as 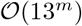, where *m* is the number of elements in the system and is unrealizable in practice for all but the smallest dynamical systems (c.f. Section 6.1).

#### Algorithm 1

Pseudocode Overview of the Routine to Calculate Phi Max

### 2.2 Example: AND+OR System

To demonstrate the problems inherent in the mathematical definition of Φ^*Max*^ we will consider a simple system comprised of an `AND` gate and an `OR` gate connected to each another, as shown in Figure 1. Since there are only two elements, we need not worry about the outermost optimization as a subsystem must be comprised of at least two components in order to generate Φ^*MIP*^ > 0, so Φ^*MIP*^ = Φ^*Max*^ in what follows. To calculate Φ^*MIP*^ we first must initialize the system into a given state. We assume an initial state *s*_0_ = 00 in all that follows, though our results are not sensitive to this choice. The next step is to identify the cause-effect structure (CES) or constellation *C* corresponding to the transition probability matrix (TPM) of the unpartitioned system. To do this, one must find the *core cause* and *core effect* of every potential mechanism in the system, where a mechanism is any element in the power set of the subsystem. In our case, the potential mechanisms are in 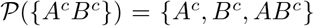 where the superscript *c* denotes the mechanism in its current state. For each element in this set, we must identify how well it constrains elements in the past power set 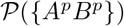, known as the past purview, as well as how well it constrains elements in the future powerset 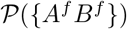, known as the future purview.

**Figure 1:**
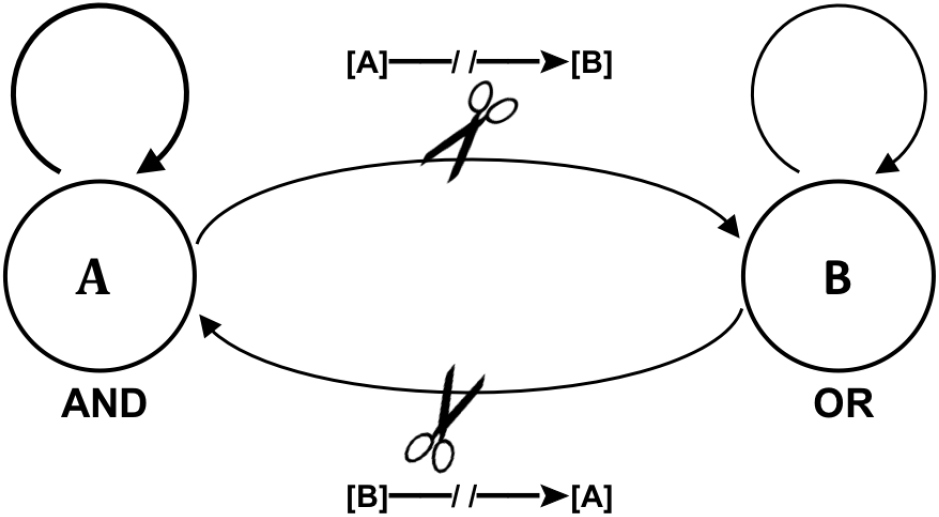
A simple system comprised of a fully connected `AND+OR` gate system. Nodes are labeled as *A* and *B*, respectively. Partitions are found by cutting the connection from one element to the other in a unidirectional fashion.

Next, we measure the earth mover’s distance *D* between the constrained distribution of each purview element and the constrained distribution of each purview element under the minimum information partition (MIP):

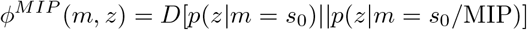

where *z* is the purview element and *m* is the mechanism. The distribution *p*(*z|m* = *s*_0_) tells us the likelihood of *z* given the current state of *m* is *s*_0_ which, compared to an unconstrained distribution, tells us how much information *m* is generating about *z*. However, we also need to know whether or not that information is “integrated” so we must break *m* and *z* up into all possible parts and ask whether the parts acting independently can generate the same amount of information as the whole. For example, to find how much integrated information is generated by the mechanism *A^c^* about the purview element *z* = *AB^p^* we calculate the probability distribution *p*(*AB^p^*|*A^c^* = 0) and compare this to the two possible partitions of the purview: *A^c^/AB^p^* → (*A^c^/A^p^* × []*/B^p^*) and *A^c^/AB^p^* → (*A^c^/B^p^* × []*/A^p^*). The first partition allows *A^c^* to constrain *A^p^* but leaves *B^p^* unconstrained (denoted by an empty bracket []) while the second partition allows *A^c^* to constrain *B^p^* but leaves *A^p^* unconstrained. The distributions generated by these partitions, shown in Figure 2, are then compared to the distribution generated by the unpartitioned system, and the partition that minimizes the earth mover’s distance to the unpartitioned system is the MIP for this purview/mechanism combination. If multiple partitions yield the the same earth mover’s distance to the unpartioned system, as is the case in Figure 2, it is irrelevant which one is chosen as all that moves forward in the computation is the scalar value of *ϕ^MIP^*.

**Figure 2:**
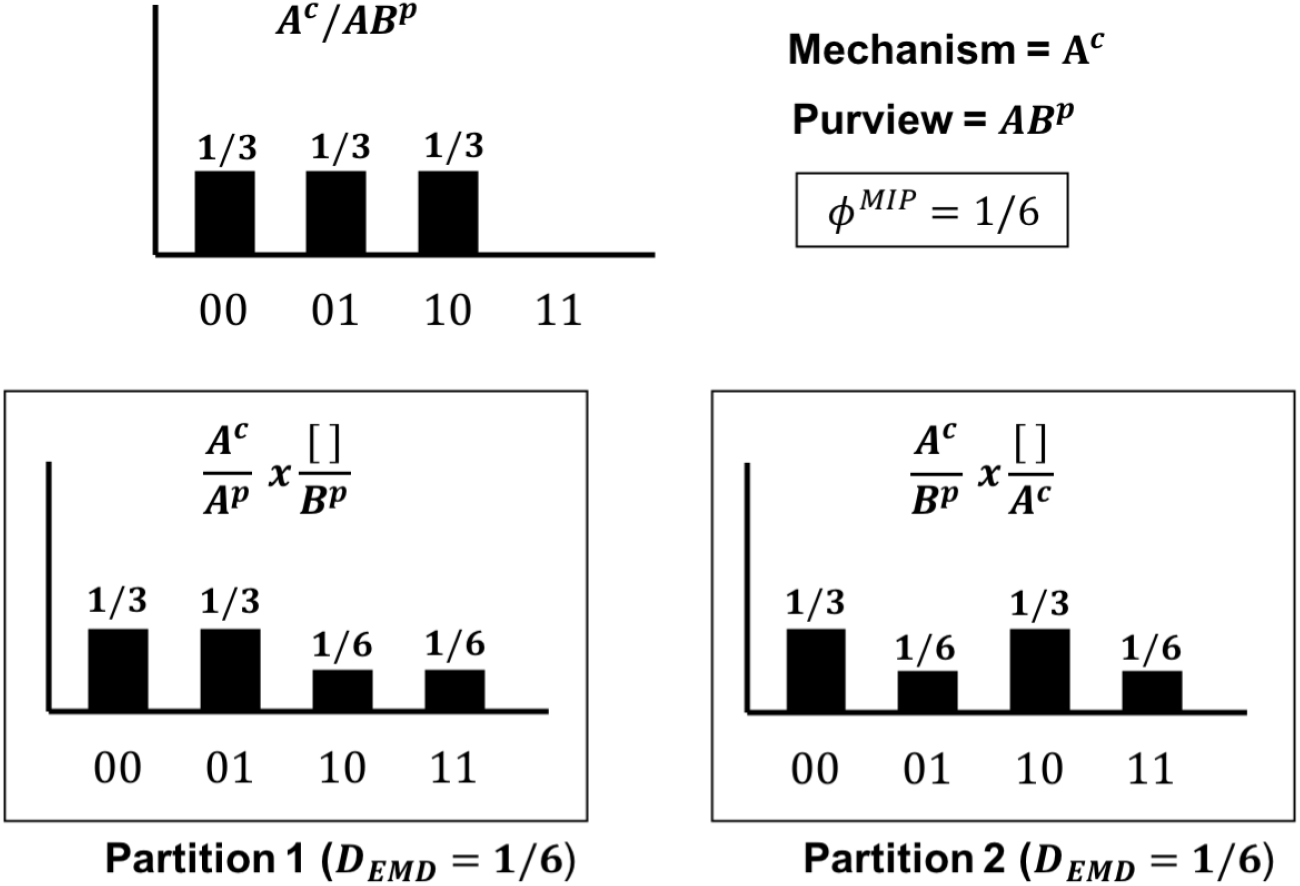
All possible partitions of a given mechanism and purview combination and the resulting *ϕ^MIP^* value.

Once we have identified the MIP (and calculated *ϕ^MIP^*) for all purview elements for a given mechanism, we define the core cause and core effect as the past and future purview elements with the greatest *ϕ^MIP^*. We denote the integrated information of the former as 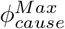 and of the latter as 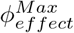 and define the total integrated information *ϕ^Max^* of a given mechanism as:

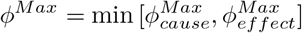

If *ϕ^Max^* > 0 for a mechanism we say that the mechanism gives rise to a concept. A concept is fully specified by three things: its *ϕ^max^* value, the cause repertoire corresponding to the core cause, and the effect repertoire corresponding to the core effect. We have already provided an example of a cause repertoire in Figure 2, namely, it is the distribution over previous states of the purview element, given the current state of the mechanism. In Figure 2, the state of *A^c^* constrains the probability of observing *AB^p^* and this constrained distribution is the cause repertoire for the purview element *AB^p^*. Any element not included as part of the purview is left unconstrained and must be independently “noised” [11]. For example, if the purview of mechanism *A^c^* is *A^p^* we generate the constrained distribution *p*(*A^p^*|*A^c^* = *s*_0_) and combine this with the unconstrained distribution for *B^p^* (denoted *p^uc^*(*B^p^*)). For the `AND+OR` system, we have *p*(*A^p^*|*A^c^* = 0) = [2/3, 1/3] and *p^uc^*(*B^p^*) = [1/2, 1/2] which yields the cause repertoire: [1/3, 1/3, 1/6, 1/6], where states are ordered in binary with the most significant digit on the left (i.e. [00, 01, 10, 11]).

The effect repertoire for a given purview element is generated in the same way as a cause repertoire. For example, the probability of observing *A^f^* given the state of mechanism *A^c^* = 0 is *P* (*A^f^*| *A^c^* = 0) = [0, 1]. Combining this with the unconstrained future distribution for *B^f^* yields the effect repertoire [1/4, 3/4, 0, 0]. Note, this is an example where the unconstrained future distribution for *B^f^* is not uniform. This is a direct result of the noising procedure: an `OR` gate receiving uniform random input is three times as likely to be in state 1 as it is to be in state 0. Furthermore, when more than one target node is involved, we must send *independent* noise to each target (to avoid correlated input). For example, in a three-node system if we were looking at the purview element *BC^f^* and noising over the state of *A^c^*, we must imagine that *A^c^* has the ability to send different signals to *B^f^* and *C^f^* at the same time (hence the term “noising”).

We can now define the CES (constellation) *C* as the set of all concepts for the system in the given state. Recall, each concept corresponds to a single mechanism and is comprised of the mechanism’s core cause repertoire, core effect repertoire, and *ϕ^Max^* value. In our example, there are at most three concepts, corresponding to the mechanisms {*A^c^, B^c^, AB^c^*}. The core cause repertoire for a mechanism is found by optimizing over all past purview elements and identifying the purview with the highest *ϕ^MIP^*, where *ϕ^MIP^* is found by further optimization over all possible partitions. Figure 3 shows *ϕ^MIP^* and the corresponding partition for all possible purview elements given the mechanism *A^c^*.

**Figure 3:**
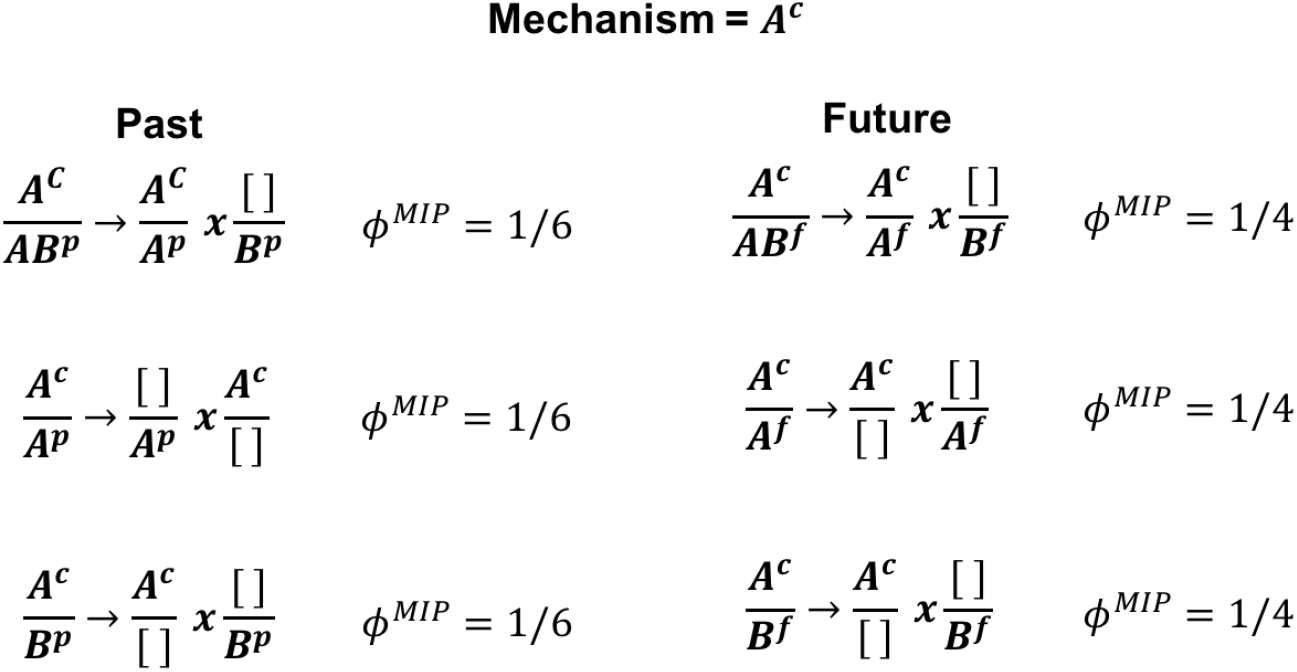
All possible purview elements and their MIPs for a given mechanism. It is here that the degeneracy is introduced, as one cannot select a unique core cause or effect for a given mechanism if there are purview element with the same *ϕ^MIP^* values.

**Figure 4:**
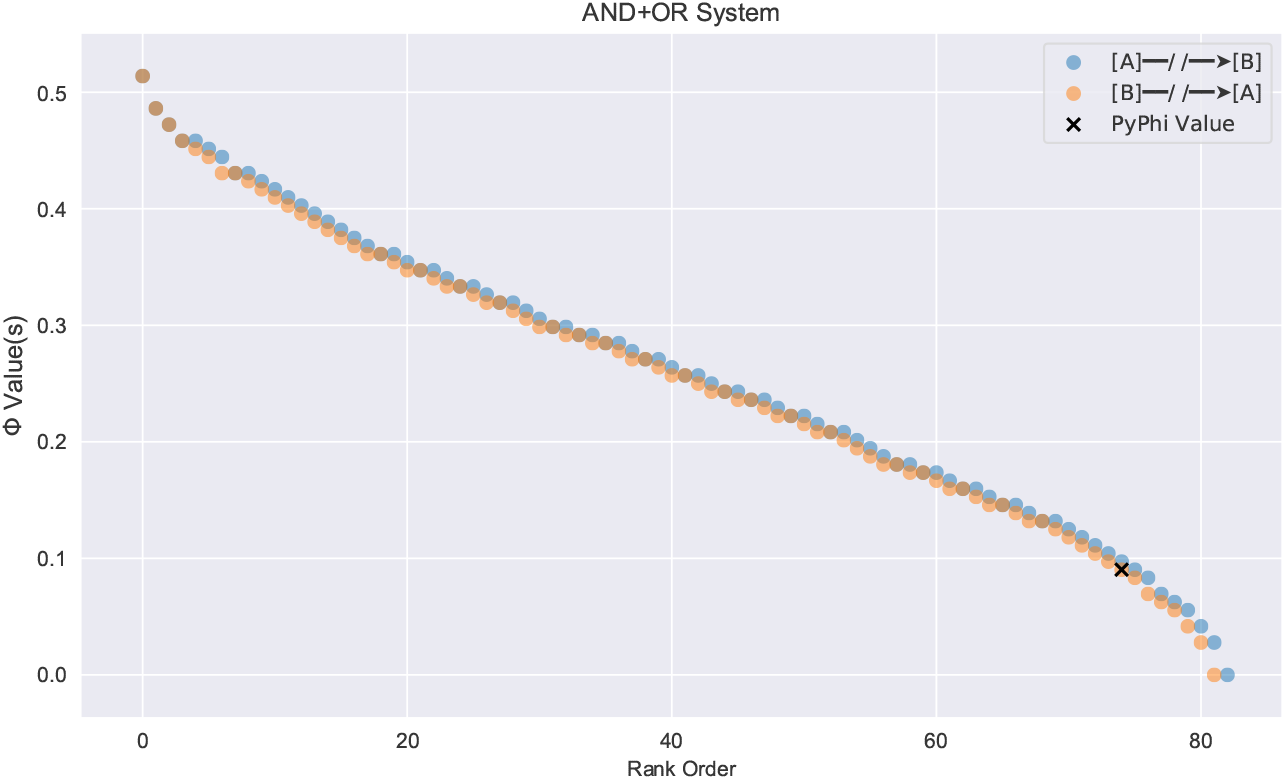
The spectrum of Φ values that result for each cut of the `AND+OR` model system. In total, there are 83 different values, all of which are equally valid Φ^*MIP*^ values according to the mathematical definition of Φ^*MIP*^. The single value corresponding to the output from the Python package `PyPhi` is shown as a black “x”.

#### 2.2.1 Degenerate Core Causes and Effects

We are now in position to see how degenerate core causes and effects can occur and their consequences. The postulates of IIT - and the exclusion postulate in particular - imply a unique core cause should be assigned to each mechanism, but the purview element that generates 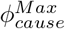 is not unique. As Figure 3 shows, *A^p^*, *B^p^*, and *AB^p^* all generate the same *ϕ^MIP^* value for the mechanism in question. Since each purview/mechanism combination is associated with a different cause repertoire, *the core cause repertoire and the resulting constellation C are not uniquely defined*. If the scalar value of 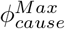 was all that mattered to the calculation of Φ, this degeneracy would be inconsequential (as is the case for partitions that generate the same *ϕ^MIP^* value for a given purview element). However, system-level integrated information Φ is defined as the cost of transforming the core cause/effect repertoires from one constellation *C* into another *C*′. That is:

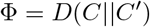

where *D* is an extension of the earth mover’s distance that calculates the cost of moving *ϕ^Max^ between repertoires*. If the core cause or effect repertoire changes, the distance between constellations will change accordingly, as the distance metric that goes into the EMD calculation is sensitive to the relative shape of the distributions and not just the scalar *ϕ^Max^* values. For example, if we were to choose *AB^p^* as the core cause as the core cause for mechanism *A^c^*, this generates the concept in *C* given by the tuple {[1/3, 1/3, 1/3, 0], [1/2, 1/2, 0, 0], 1/6} where the first element is the core cause repertoire, the second element is the core effect repertoire, and the third element is the *ϕ^Max^* value. However we could just as easily have chosen *A^p^* as our core cause and *A^f^* as our core effect. In which case, the concept generated for *A^c^* would be {[1/3, 1/3, 1/6, 1/6], [1/4, 3/4, 0, 0], 1/6}. Clearly, these choices have the same Φ^*Max*^ value but significantly different core cause and effect repertoires.

To illustrate the consequences of this, let *C* be the constellation consisting only of the concept generated by *A^c^* with core cause cause *AB^p^* and core effect *AB^f^* and let *C*′ be the constellation consisting of only the null concept for this system (the unconstrained cause and effect repertoires):

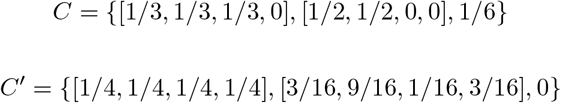

The extended earth mover’s distance is the cost of transforming *C* into *C*′ by moving *ϕ^Max^* = 1/6 a distance given by the sum of the (regular) earth mover’s distance between cause repertoires and effect repertoires. Namely, we have

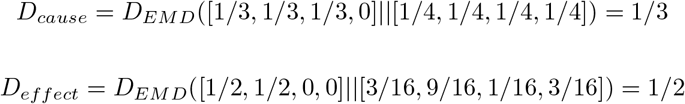

which results in the integrated conceptual information:

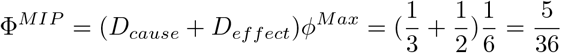

Now, if we instead choose *A^p^* and *A^f^* as our core cause and core effect we have:

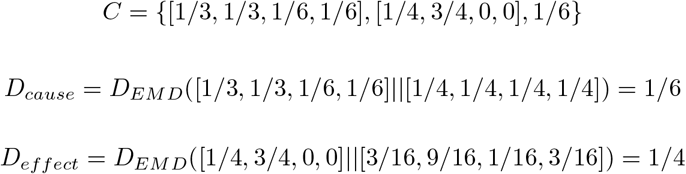

corresponding to an integrated conceptual information:

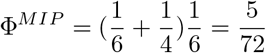

Thus, we get different values of Φ^*MIP*^ depending on our choice of core cause and core effect.

#### 2.2.2 A Spectrum of Non-unique Φ Values

Each combination of degenerate core cause/effect repertoires results in the potential for a different Φ value. For example, if the unpartitioned system has three degenerate core causes for *A^c^* and two for *B^c^*, then there are 3 × 2 = 6 non-unique constellations (CES) for the unpartitioned system. If the partitioned system also has six non-unique CES, then there are a total of 36 different combinations for the distance between constellations (Φ). For the `AND+OR` system, the unpartitioned system has a total of 81 non-unique combinations, while each cut has a total of 9 non-unique constellations. Thus, one must examine the distance between 81 × 9 × 2 different combinations of constellations in order to determine all possible Φ values, which we refer to as the “spectrum” of Φ values for the subsystem. Note, not all Φ values are valid Φ^*MIP*^ values; it is only those between the upper and lower Φ value of the minimum information partition (MIP) that satisfy the definition of Φ^*MIP*^. In total, 83 non-unique Φ^*MIP*^ values result for the `AND+OR` system, as shown in Figure 2.2.2. Again, we emphasize there is nothing in the axioms or postulates of IIT to suggest which of these 83 values IIT actually predicts, as all are equally valid according to the mathematical definition of Φ. Crucially, both Φ^*MIP*^ = 0 and Φ^*MIP*^ > 0 are present in the spectrum, meaning IIT cannot actually predict whether or not this simple system is conscious (or conversely, one could interpret the theory to simultaneously predict the system is both conscious and unconscious).

## 3 Results

We now apply our methodology to a variety of recently published systems where Φ was calculated, in order to determine the extent to which degenerate core causes/effects lead to a need for revised interpretation of reports that provide only a unique Φ value. We begin with a pedagogical case study, followed by the broad application of the `PyPhi-Spectrum` package, described in Section 6.3, to as large of a corpus as is computationally feasible.

### 3.1 Case Study: Three-node Fission Yeast Cell Cycle

As a case study, we consider the Boolean network model of the fission yeast cell cycle from Marshall et al. [16]. In this study, IIT is used to analyze the causal structure of a minimal biological system, namely, the cell cycle of the fission yeast *S. pombe*. Using Φ, the authors identify three integrated subsystems corresponding to the full system (eight nodes), a six node subsystem, and a three-node subsystem - all potentially of biological importance. Of these three systems, only the smallest may be subject to our analysis, though we expect similar results for the other two systems. Applying the methodology from Section 2, we find a spectrum of *244 non-unique* Φ *values*, spanning a range from 0.00 – 0.83 bits, see Figure 5. Crucially, only one of these values (Φ = 0.09) is published as the unique Φ^*MIP*^ value for this subsystem. In reality, all of these values are valid according to the mathematical definition of Φ. Furthermore, the inclusion of Φ = 0 in the spectrum of possibilities changes the biological interpretation of the results entirely. If the subsystem under consideration has Φ^*MIP*^ = 0, rather than Φ^*MIP*^ > 0, it would not be identified as “integrated” and its biological function would not be deemed of interest. Thus, the conclusion that this subsystem is of biological importance is entirely dependent on the arbitrary selection of a single Φ value from the spectrum of possibilities and, in general, it is impossible to tell *a priori* whether or not the narrative being built around a particular Φ value is valid without studying the entire spectrum of possibilities.

**Figure 5:**
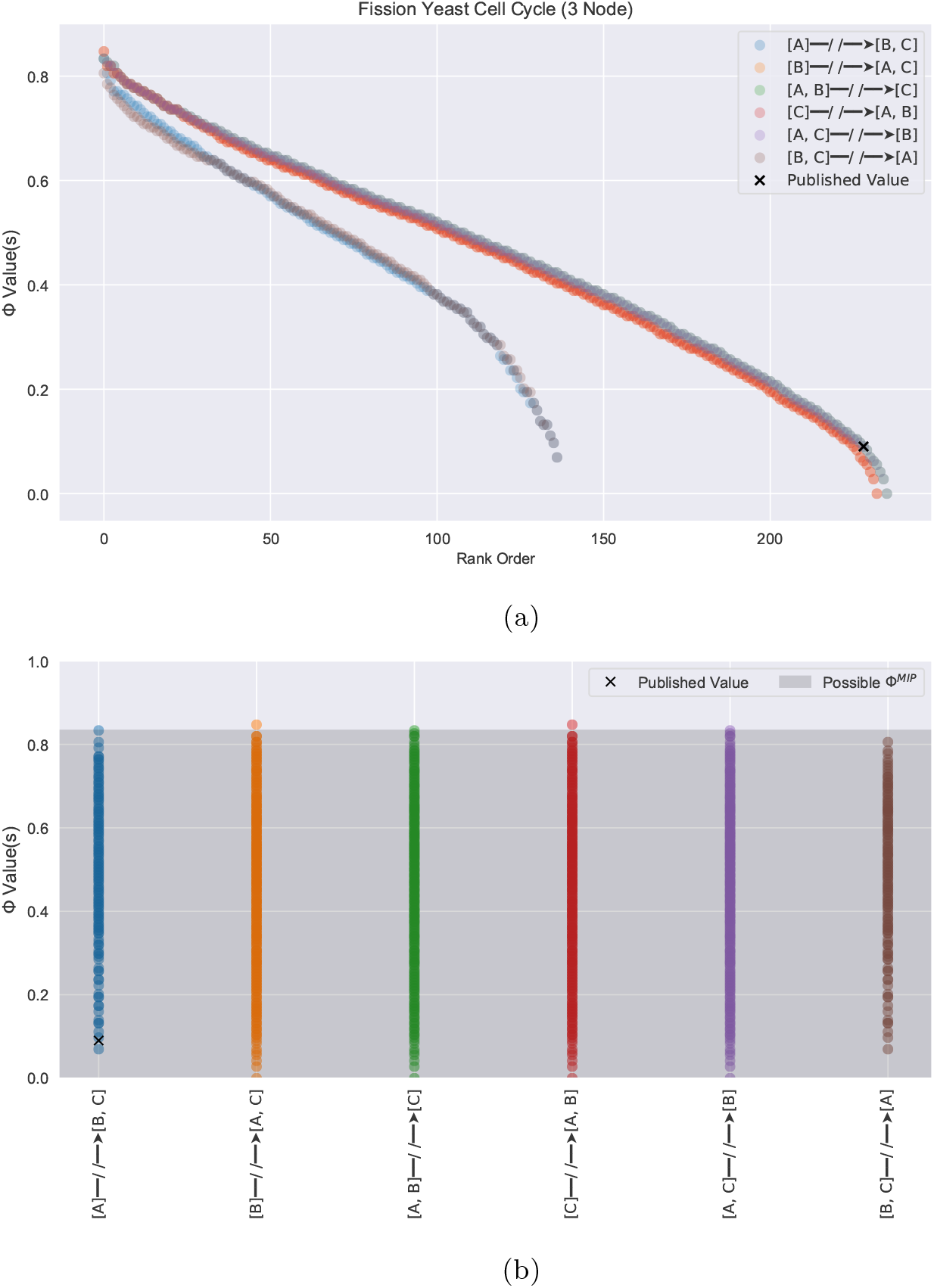
The spectrum of Φ^*MIP*^ values that result from degenerate core causes/effects in the three-node fission yeast system analyzed by Marshall et al [16]. The published value for this system is shown as a black “x”. Figure (a) shows the Φ values for each cut in rank order, while (b) shows the Φ values for each cut relative to the upper and lower bound on Φ^*MIP*^. All Φ values between the min and max Φ value of the MIP are equally valid Φ^*MIP*^ values.

### 3.2 The Non-uniqueness of Published Φ Values

Next, we use `PyPhi-Spectrum` to anaylze a corpus of recently published Φ values, with the goal of understanding the extent to which underdetermined qualia affect interpretation of previously published Φ. The corpus we analyze is intended to be comprehensive, but we are limited by the computational resources required to perform these calculations (see Section 6.1). Of the dozen or so Φ values that are published in the literature [16–19, 21–26, 28–32], only a handful are small enough to be subjected to our analysis [11, 16, 18, 21, 22, 24, 26, 32]. These systems, summarized in Table 1, are selected primarily for their size, though size alone is not a good indication of computational tractability. For example, certain three-node systems, such as that found in Hanson and Walker 2020 [24], have thousands of degenerate cause-effect structures while others, such as that found in Farnsworth 2021 [18], have just a few. It is not readily apparent what dictates the number of non-unique cause-effect structures that result for a given system, though symmetric inputs almost certainly play a role (see Section 4). Consequently, our corpus is limited to small systems (2-4 nodes) which happen to allow relatively fast evaluation via `PyPhi-Spectrum`^2^. While improvements such as parallelization could certainly be made to improve performance and increase the size of our corpus, we do not believe doing so would add much to the interpretation of our results.

**Table 1:**
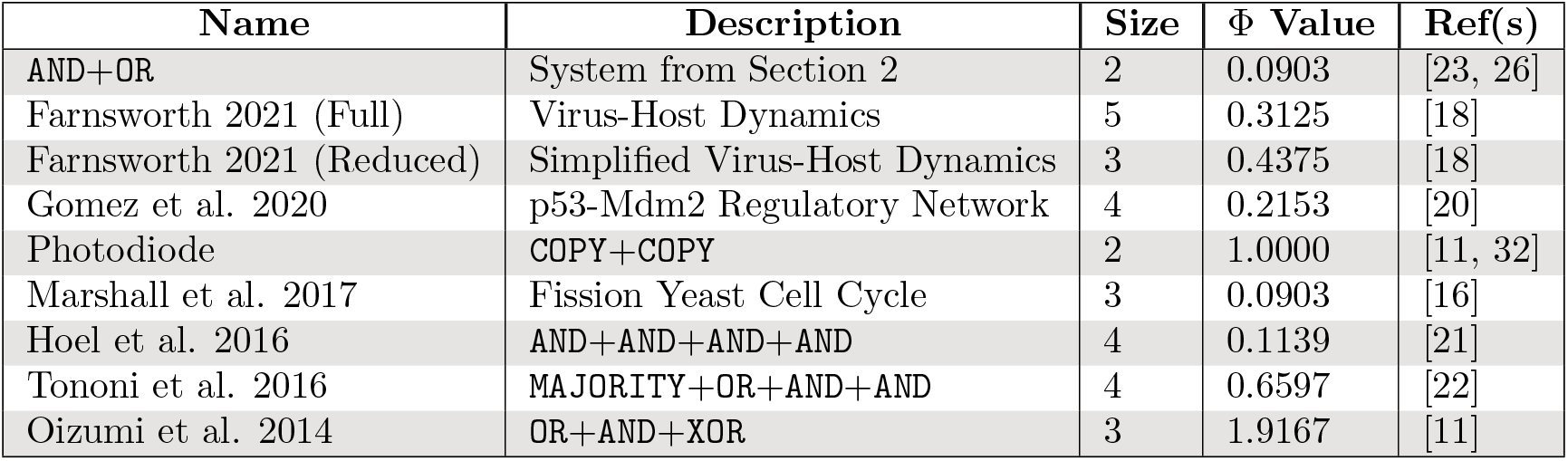
Summary of corpus ordered reverse chronologically. Sources were selected based on the publication of a unique Φ^*MIP*^ value and computational tractability. Additional details required for analysis, such as transition probability matrices and initial states are provided in Section 6.4.

| Name | Description | Size | $\Phi$ Value | Ref(s) |
| --- | --- | --- | --- | --- |
| AND+OR | System from Section 2 | 2 | 0.0903 | [23, 26] |
| Farnsworth 2021 (Full) | Virus-Host Dynamics | 5 | 0.3125 | [18] |
| Farnsworth 2021 (Reduced) | Simplified Virus-Host Dynamics | 3 | 0.4375 | [18] |
| Gomez et al. 2020 | p53-Mdm2 Regulatory Network | 4 | 0.2153 | [20] |
| Photodiode | COPY+COPY | 2 | 1.0000 | [11, 32] |
| Marshall et al. 2017 | Fission Yeast Cell Cycle | 3 | 0.0903 | [16] |
| Hoel et al. 2016 | AND+AND+AND+AND | 4 | 0.1139 | [21] |
| Tononi et al. 2016 | MAJORITY+OR+AND+AND | 4 | 0.6597 | [22] |
| Oizumi et al. 2014 | OR+AND+XOR | 3 | 1.9167 | [11] |

Our primary result is shown in Figure 6, which reports calculated values for the entire spectrum of Φ^*MIP*^ values relative to the published value for every text in our corpus. There are several things to note. First, the existence of a unique Φ value is rare: only the photodiode has a spectrum consisting of a single value (the number of different Φ^*MIP*^ values is denoted by |Φ^*MIP*^| in Figure 6). For the rest of the corpus, the spectra often consist of dozens if not hundreds of non-unique Φ^*MIP*^ values, of which only one is published (denoted as a black “*x*” in Figure 6). Importantly, we do find cases where the spectrum contains both Φ = 0 and Φ > 0 values, indicating predictions consistent with IIT that allow interpretation of a system as conscious and unconscious depending on the value chosen. This occurs for three out of the ten published Φ values in our corpus: `AND+OR`, Marshall et al. 2017 [16], and Hoel et al. 2016 [21]. This has significant implications for the logical foundations of the theory (the exclusion postulate in particular). Additionally, the span of Φ^*MIP*^ values is often comparable to the entire range of possibilities expected for systems of this size. A typical Φ spectrum spans roughly 1/2 of a bit (Figure 6). In comparison, a (deterministic) two-node Boolean system is bounded from above by Φ^*MIP*^ ≤ 1.5 bits (Section 6.2), which implies that the Φ values calculated by IIT are not only non-*unique* but also non-*specific* (i.e. they don’t constrain the possible Φ values to a small portion of the range).

**Figure 6:**
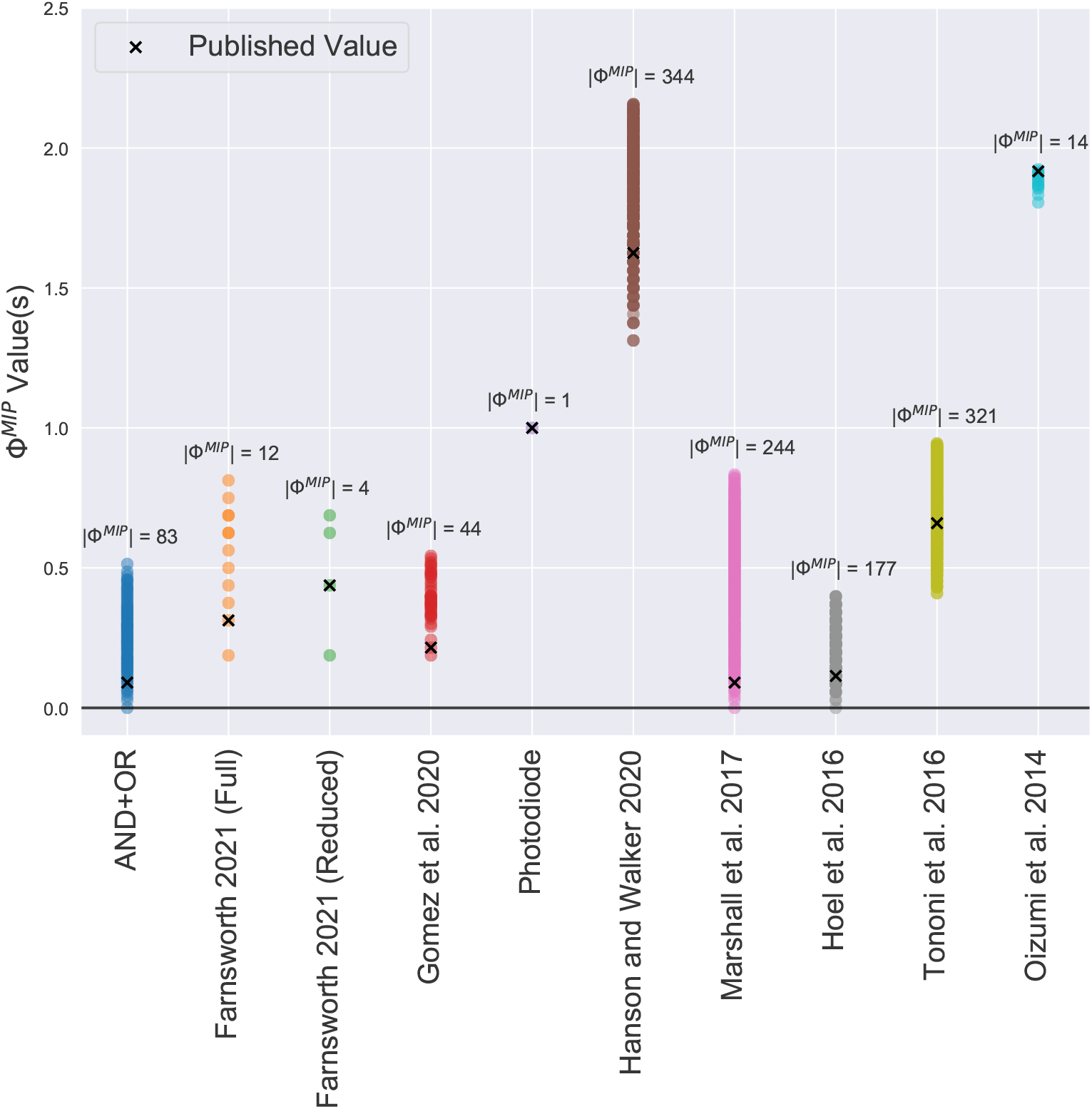
Possible Φ^*MIP*^ values relative to the published value for the corpus shown in Table 1. Each point represents a possible Φ^*MIP*^ value that results from the mathematical definition of Φ. The number of different Φ values for each system is given by the cardinality of its spectrum |Φ^*MIP*^|.

## 4 Existing Solutions

The problem of degenerate core causes/effects in IIT is already recognized, but so far its impact has not be extensively studied and therefore there has not been sufficient motivation to address it [11, 13, 14, 26]. Previous studies have pointed to the problem of degenerate core causes/effects in IIT (although not studying the extensive impact on reported literature as we do here). To our knowledge, there are several different solutions to this problem, with differing degrees of justification, which we review next.

The first solution is that put forward by Oizumi et al. in IIT 3.0 [11]. In Figure S1 of their Supporting Information, the authors argue that the degenerate core cause corresponding to the *biggest purview* element should be selected as the core cause. The justification is that the larger purview “specifies information about more system elements for the same value of irreducibility”: that is, the *ϕ^max^* values of the degenerate purview elements are the same (same value of irreducibility) but bigger purview elements constrain more of the system (e.g. *AB^p^* constrains more of the system than *A^p^*). This motivation however does not make direct connection to the axioms and/or postulates of the theory [14], meaning another choice could be equally consistent with the phenomenology of the theory. Additional postulates are therefore required. The degenerate core causes in the example they consider correspond to purview elements of different sizes, which allows this simple criterion to result in a unique core cause. However, we have already shown that in general there is no guarantee this is the case. Therefore, this criterion cannot be used to guarantee a unique Φ value and additional criterion would need to be added..

By contrast, Krohn and Ostwald (hereafter KO17) reach the exact opposite criterion as a proposed solution to the same problem [13]; namely, KO17 argue that it is the *smallest purview* element that should be selected as the core cause/effect in the case of tied purviews. Their motivation is “causes should not be multiplied beyond necessity” [12]. A similar language is found in the phenomenology of IIT in the exclusion postulate, but it is not clear that it can be applied to the dimensionality of purview elements. Further motivation is required to illustrate why choosing *B^p^* as the core cause of *A^c^* instead of *AB^p^* multiplies the cause of *A^c^* beyond necessity. Additionally, again we encounter that the purview element is not guaranteed to be unique, which means Φ is still not mathematically well-defined. To address this, KO17 present a completely novel solution in the form of a modified definition of Φ based on the difference in the sum of *ϕ^max^* values between constellations, rather than the extended earth mover’s distance. The benefit of this definition is that it depends only on the scalar *ϕ^max^* values associated with concepts, rather than the non-unique cause/effect distributions. In other words, it does not matter which of the degenerate core causes is selected because they all have the same *ϕ^Max^* value and the integrated conceptual information is just the sum of *ϕ^Max^* values over concepts.

Another solution is the “differences that make a difference” criterion proposed by Moon [14]. Here, the author argues that if degenerate core causes/effects exist then *none* of the corresponding purview elements should be selected as the core cause/effect (i.e. *ϕ^max^* = 0 for the mechanism). This solution is based on the idea that in IIT “to exist is to cause differences”. If tied purview elements exist with the same *ϕ^Max^* value, then the *ϕ^Max^* value does not change if one of the tied purview elements is excluded. For example, if *A^p^* and *B^p^* both give rise to a concept with *ϕ^Max^* = 1/6, one can eliminate *B^p^* without changing the *ϕ^Max^* value for the mechanism; therefore, the existence of *B^p^* does not make a difference “from the intrinsic perspective of the system”. The fact that one can do this individually for each of the degenerate core causes or effects implies that none can give rise to a concept and *ϕ^Max^* = 0 for the mechanism.

Problems arise with the Φ values that result from all four proposed solutions to the non-uniqueness problem. Namely, selecting the smallest or largest purview element requires additional phenomenological assumptions, and even with these would not guarantee a unique Φ value. The KO17 and Moon criteria result in Φ = 0 for systems that are clearly integrated. In the case of the KO17 definition of Φ, under a partition, the sum of *ϕ^Max^* values can actually *increase*, resulting in the confounding conclusion that the system is integrating more information after a cut than it was before (Φ < 0). These so-called “magic cuts” are discussed by Krohn and Ostwald [13], as well as in the PyPhi documentation. More immediately, if we apply the KO17 definition of Φ to the `AND+OR` system from Section 2, we find Φ = 0 due to the fact that the *ϕ^Max^* values for each concept are the same before and after the system level minimum information partition (Σ*ϕ^Max^* = 5/12 in both cases). Consequently, we reject this modified definition of Φ outright based on the idea that an `AND+OR` system *is* integrated, *e.g.* it satisfies the general mathematical definition of integrated information, which is an inability to tensor factorize a system without changing the underlying dynamics [33, 34]. In the case of the `AND+OR` system, one cannot cut *A* from *B* without affecting the state of *B*, and vise versa, which implies the system is integrated. The same conclusion applies to Moon’s criterion, for which the `AND+OR` system again fails to yield Φ > 0 due to the presence of degenerate core causes/effects for all mechanisms. In addition, the “differences that make a difference” argument relies entirely on the assumption that the *ϕ* value being measured is an accurate reflection of what it means to make a difference. Put simply, cutting information from an `AND` gate to an `OR` gate *does make a difference* in terms of system-level integration, and the only way to justify that it doesn’t is to define differences that make a difference in terms of *ϕ* values.

As demonstration of these problems inherent with each of the four existing solutions, we reanalyze our corpus enforcing each criterion in turn (Figure 7)^3^. As expected, the KO17 and Moon solution yield Φ = 0 for several systems that are integrated (*e.g.*, that cannot be tensor factorized without changing the underlying dynamics). The “smallest” criterion does little to mitigate the degeneracy. At first glance, however, it appears the “biggest” criterion avoids both of these issues and provides a positive Φ value in all cases. This is an idiosyncrasy of our data set, as selecting the biggest purview element suffers from the same problem as selecting the smallest purview element; namely, degenerate core causes/effects are often the same size. The reason that the “biggest” solution appears to yield unique Φ values for the systems under consideration is due entirely to the ubiquitous use of two-input logic gates (`AND, OR,XOR, NOR`, etc.) in our corpus. In such cases, the tied purview elements are almost always *A*, *B* and *AB* for which selecting the largest purview element (*AB*) results in a unique core cause/effect. However, this does not hold in general, as systems comprised of logic gates with more than two inputs (e.g. neurons in the human brain) have entirely different symmetries. As Figure 8 shows, a simple system of majority gates, each with three inputs, is enough to prove that a unique Φ value does not result from the “biggest” criterion. Thus, the fundamental problem remains unaddressed.

**Figure 7:**
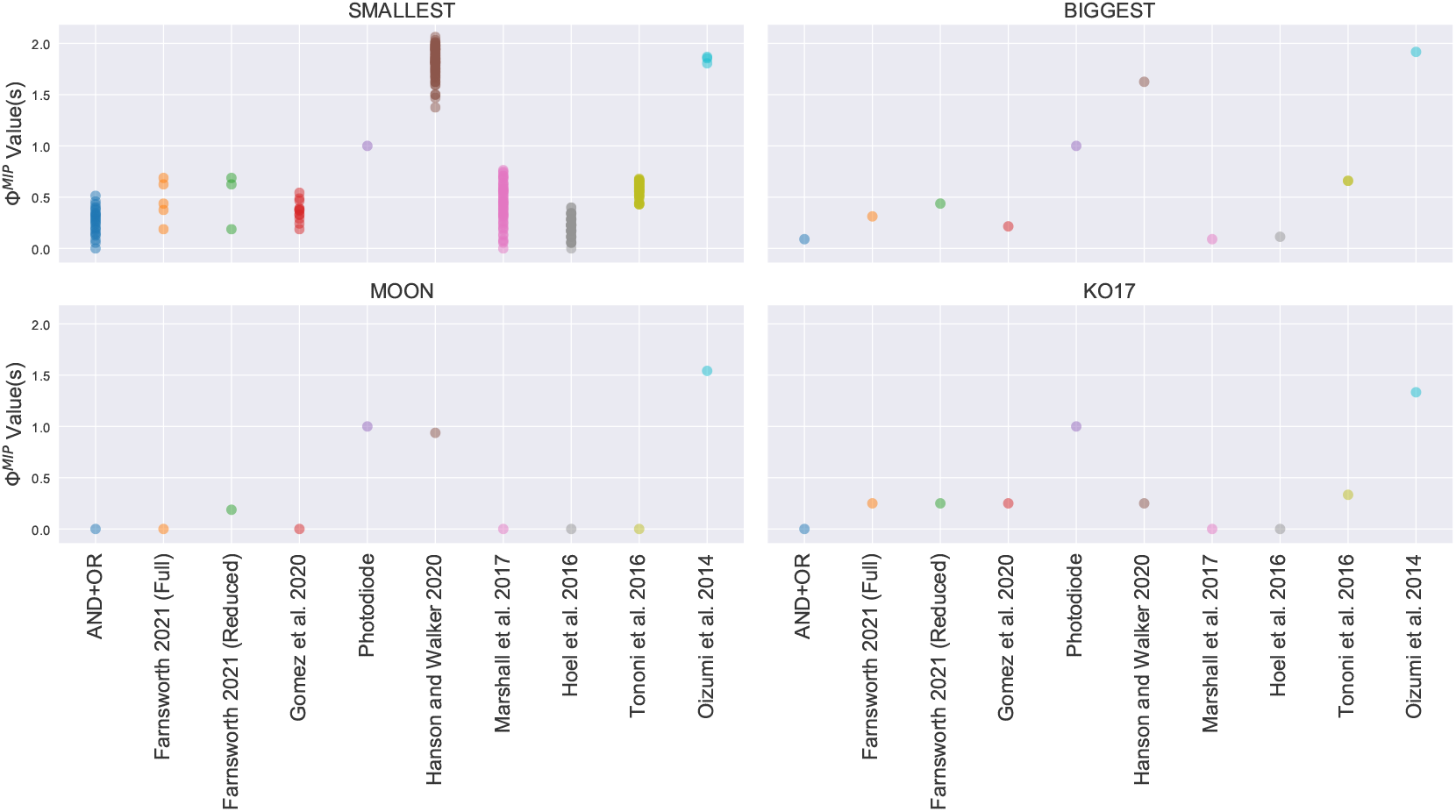
The spectrum of Φ values that results from each of the four proposed solutions. The “smallest” and “biggest” solutions do not guarantee a unique Φ value, while the Moon and KO17 solutions result in Φ = 0 for systems that are clearly integrated, such as the `AND+OR` gate.

**Figure 8:**
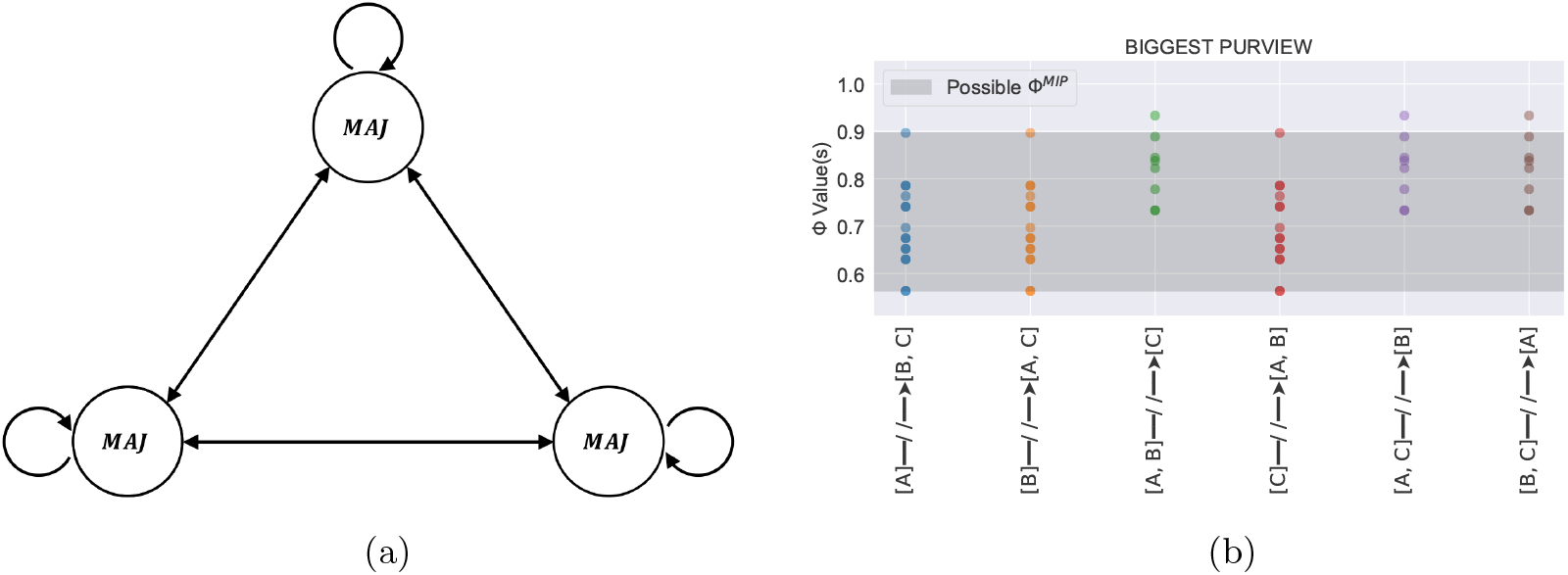
For a system of three fully connected majority gates (a) selecting the largest purview element does little to mitigate degeneracy, as evident by the range of possible Φ^*MIP*^ values (b).

## 5 Discussion

In IIT, the constellation corresponding to the unpartitioned CES is equated with nothing less than subjective experience itself. The geometric shape of this constellation (meaning the shape of the probability distributions that comprise the core cause/effect repertoires) is identified as “what it is like” to be something (c.f. [35]) while the Φ^*Max*^ value is identified as the overall “level” of consciousness. Our results demonstrate how this structure is underspecified. If held to IIT’s own criteria, Φ cannot be used to measure the contents or quantity of subjective experience. In the words of Moon, “the qualia underdetermination problem shakes IIT to the ground” [14].

It is critical that such challenges be addressed, especially given how IIT is more popular than ever and Φ is being used to bolster arguments about the nature of consciousness in neuroscience and beyond [17, 18, 36]. While it is important to acknowledge that IIT is not necessary tied to its current mathematical formalism, it is equally important to point out that the theory demonstrates several hallmark features of an unscientific handling of ideas to date [37]. The first is the use of an ad hoc procedure to resolve fundamental contradictions inherent in the theory: IIT tries to resolve the undetermined qualia problem with an ad hoc procedure of selecting the smallest or largest purview element. The second is a lack of transparency with regard to the mathematical implementation of the theory: `PyPhi` essentially acts as a black-box algorithm that buries many of the mathematical problems inherent in the Φ calculation, most notably the arbitrary selection of a unique Φ value. And third, the theory struggles to ground itself experimentally: recent proofs have demonstrated that IIT is either falsified or inherently unfalsifiable depending on the inference procedure that is used to independently test predictions from the theory. In combination, these three features strongly suggest that IIT is on the wrong side of the demarcation problem.

Granted, there is hope that IIT may yet provide a blueprint for a successful scientific theory of consciousness [38, 39]. Culturally, however, what we see is that IIT is presented as a self-consistent theory, without resolving contradictions inherent in its logical foundation that imply it is in fact not self-consistent [14, 40, 41]. On one hand, this could be seen as a positive, as Feyerabend argues that all new theories must proceed counterinductively for a matter of time if they are to succeed. That is, they should go against well-established principles and hold onto assumptions in the face of overwhelming evidence to the contrary [42]. Yet, even Feyerabend is careful to distinguish between scientific and unscientific handling of ideas, stating:

> The distinction between the crank and the respectable thinker lies in the research that is done once a certain point of view is adopted. The crank usually is content with defending the point of view in its original, undeveloped, metaphysical form, and he is not prepared to test its usefulness in all those cases which seem to favor the opponent, or even admit that there exists a problem. It is this further investigation, the details of it, the knowledge of the difficulties, of the general state of knowledge, the recognition of objections, which distinguishes the “respectable thinker” from the crank. The original content of his theory does not. [43]

Going forward, it is of paramount importance that proponents of IIT seek to address the difficulties associated with such problems head-on, rather than bypassing them, as the scientific merit of the theory will ultimately depend on it. This consideration is especially important given the negative repercussions that could result from the misapplication of a theory of consciousness in medical, legal, and moral settings.

## 6 Appendix

## 6.1 On the Computational Complexity of Φ

For a subsystem of size *n*, the computational complexity scales as follows. First, one must calculate the cause-effect structure (CES) for every possible partition of the subsystem. If the partition is a bipartition, as is typically assumed [13, 15], the number of ways to do this is *S*(*n,* 2), where *n* is the size of the subsystem and *S*(*n, m*) are Stirling numbers of the second kind [44]. However, two small corrections should be considered: first, partitions are unidirectional, and second, the unpartitioned system must also be addressed. The former consideration results in twice as many bipartitions, while the latter results in a single additional partition. Combining these results in a total of 2*S*(*n,* 2) + 1 partitions which, for large *n*, is well approximated as 2*S*(*n,* 2). For each CES, there are 2^*n*^ − 1 potential mechanisms, corresponding to the size of the powerset of elements excluding the empty set. For each potential mechanism, there are 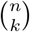 purview elements of size *k*, each of which can be partitioned *S*(*k*, 2) times. Therefore, there are 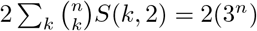 elementary distance calculations that must be performed to calculate a single CES, where the additional factor of two is due to the need to optimize *ϕ^max^* over both past and future purviews. Putting this together, there are a total of 2*S*(*n,* 2 + 1) × (2^*n*^ − 1) × 2(3^*n*^) ≈ 12^*n*^ elementary distance calculations required to get the system-level integrated information Φ^*MIP*^ for a given subsystem. For the global system, this calculation must be embedded in an additional optimization corresponding to maximizing over the powerset of all possible subsystems (i.e. Φ^*Max*^ = max{Φ^*MIP*^}). For a global system of size *m*, there are 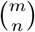 subsystems of size *n*, each with 12^*n*^ elementary distance calculations. Therefore, there are a total of 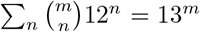 elementary calculations required to find Φ^*max*^ for a global system of size *m*. For all but the smallest *m* values, the computational resources required to actually calculate Φ^*Max*^ are impossible to realize.

Interestingly, the 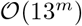 scaling derived here is in tension with the previously published value of 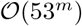 [15]. This could be due to the possibility that the 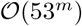 scaling considers all possible partitions, rather than strict bipartitions, or perhaps it resolves the elementary computation in terms of some more fundamental operation (e.g. bit flips). Without additional information, it is difficult to say whether or not either of these considerations could resolve the tension between values. We do note, however, that the published values of *t* = 1, *t* = 16, and *t* = 9900*s* for *n* = 3,*n* = 5, and *n* = 7 ([15]) are within an order of magnitude of the predicted 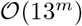 scaling while off by 40 orders of magnitude from an 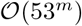 scaling, though the use of parallelization complicates this point.

## 6.2 Calculating an Upper Bound on Φ

It is relatively straightforward to calculate a loose upper bound on Φ^*MIP*^ for a subsystem of size *n*. To do so, one need only understand the extension of the earth mover’s distance *D* that is used in the calculation Φ^*MIP*^ = *D*(*C*||*C*_→_). By definition, the “earth” being moved is *ϕ^Max^* between concepts in the unpartitioned CES (C) and the partitioned CES (*C*_→_), while the “distance” it is moved is measured by the regular earth mover’s distance between concepts [11]. In light of this, a straightforward upper bound on Φ^*MIP*^ can be found by asking what the maximum value of *ϕ^Max^* is for each concept and moving that amount as far away as possible. For a mechanism of size *m*, its *ϕ^Max^* value is bounded from above by the maximum value of the regular earth mover’s distance, which is *EMD^Max^*(*m*) = *m*. It is easy to see this is the case, as *EMD^Max^* is achieved when all the probability (*p* = 1) is moved the maximum Hamming distance (*H^Max^*), which is *m* for a mechanism comprised of *m* bits. For example, *EMD^Max^* for a three-bit mechanism is achieved when *p* = 1 is moved from state 000 to state 111 (*H^Max^* = 3), so *ϕ^max^* = *EMD^Max^* = 3. Next, we must ask what the maximum distance *D^Max^* in conceptual space is that this amount of *ϕ^Max^* can be moved. Since this distance is again a regular earth mover’s distance, we have *D^Max^* = *EMD^Max^*(*m*) = *m*. Thus, the maximum contribution a mechanism of size *m* can make to the extended earth mover’s distance *D* is upper bounded by *ϕ^Max^*(*m*)*D^Max^*(*m*) = [*EMD^Max^*(*m*)]^2^ = *m*^2^. Of course, not all mechanisms are the same size, so the total contribution is bounded by the sum of the maximum contribution from mechanisms of each size, namely:

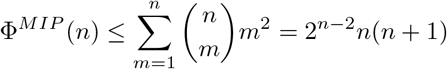

To date, this is the only known upper bound on Φ^*Max*^ that we are aware of (though bounds on *ϕ^max^* and IIT 2.0 are readily available [13, 34, 45–47]), and it is a very loose bound. For a subsystem of size *n* = 2, as is the case for the `AND+OR` system we consider in the main text, we have Φ^*MIP*^ (2) ≤ (1)^2^ +(1)^2^ +(2)^2^ = 6 bits. In practice, we cannot reasonably expect *ϕ^Max^* = *EMD^Max^* for all mechanisms, as the existence of *ϕ^Max^* = *EMD^max^* for one mechanism almost certainly precludes the existence of *ϕ^Max^* = *EMD^Max^* for another. Likewise, cutting a CES cannot possibly result in a distance of *D^Max^* = *EMD^Max^* for all concepts, as additional noise cannot be used to increase the fidelity of constraints. At best, it is likely that concepts map to the null concept in the *CES* of the MIP, corresponding to a maximum distance *D^Max^*(*n*) = *n/*2. In this case, the bound that results is Φ^*MIP*^ ≤ 2^*n*−3^*n*(*n* + 1), which is still likely loose. To tighten it, one must consider the *ϕ^Max^* values that can result for a system of mechanisms as an ensemble, rather than individually, which is a task that we found quickly became intractable.

## 6.2.1 Numerical Approach

Fortunately, for our purposes a numerical approach will suffice. Given a small enough system, it is possible to calculate the Φ values for every possible transition probability matrix (TPM) that results from Boolean logic on a two-bit system. Namely, each bit (*A* and *B*) takes one of two possible states in response to the global state of the system. This means there are 2^4^ = 16 possible state transitions for each coordinate, for a total of 16^2^ unique TPMs. For each, it is possible to calculate the Φ spectrum that results using the algorithm we describe in the main text. Then, the upper bound on Φ^*MIP*^ is simply the maximum Φ^*MIP*^ value over all possible TPMs in all possible initial states. Since the system is only two bits, this bound on Φ^*MIP*^ is equivalent to the bound on Φ^*Max*^, as subsystems must be comprised of at least two bits to generate Φ^*MIP*^ > 0. Performing this exercise results in the bound Φ^*Max*^ ≤ 1.5, which is interestingly exactly 1/4 the analytical bound derived in the previous section; as discussed, it is likely that a factor of 1/2 is accounted for if *D^Max^*(*n*) = *n/*2, while the other factor of 1/2 may be accounted for by the same type of argument applied to *ϕ^Max^* (rather than *D^Max^*). If so, the upper bound on Φ^*MIP*^ would be 2^*n*−4^*n*(*n* + 1) and is potentially more tractable to calculate than previously believed.

## 6.3 The PyPhi-Spectrum Package

The Python package PyPhi (available for download at http://integratedinformationtheory.org) provides all of the basic functionality needed to calculate the spectrum of Φ values for a given system. Namely, it allows the user to calculate purviews, cause/effect repertoires, earth mover’s distance, CES distance (extended earth mover’s distance), and more. In addition, it contains prebuilt classes for data structures such as concepts that are useful in the calculation. Here, we wrap this basic functionality into a modified version of `PyPhi` called `PyPhi-Spectrum` that allows the user to calculate all Φ values for a given subsystem with a single function call. To install this package, one can simply download or clone the entire `Phi-Spectrum` repository (which includes core `PyPhi` functionality) from https://github.com/elife-asu/pyphi-spectrum.

A pseudo-code overview of the `PyPhi-Spectrum` wrapper is shown in Algorithm 2, while basic usage is shown in Algorithm 3. Note, the `get-phi-spectrum` call returns the Φ values that result from all possible concepts for each cut, while the `get-phi-MIP` call returns the Φ values corresponding to the minimum information partition (i.e. Φ^*MIP*^). Optimizing the latter over all possible subsystems would provide Φ^*Max*^ for a given system. To install the code, download or clone the entire `PyPhi-Spectrum` repository (which includes core `PyPhi` functionality) from https://github.com/elife-asu/pyphi-spectrum.

### Algorithm 2

Pseudocode overview of the `PyPhi-Spectrum` wrapper

### Algorithm 3

Basic Usage for the `PyPhi-Spectrum` Wrapper

## 6.4 Additional Details Related to the Calculation of Φ Values

In this section, we provide the transition probability matrices and initial states necessary to replicate our results. The same data can be found in downloadable form via the GitHub repository: https://github.com/elife-asu/pyphi-spectrum.

## 6.4.1 Photodiode [11, 32]

A photodiode is a simple system of two interacting `COPY` gates, taking input from one another. It is arguably the simplest “integrated” system one can study, and has been studied in the context of IIT at least twice [11, 32]. Following Chalmers and McQueen [32], we set the initial state of the system to be *s*_0_ = 10. The transition probability matrix is given below.

**Table 2:**
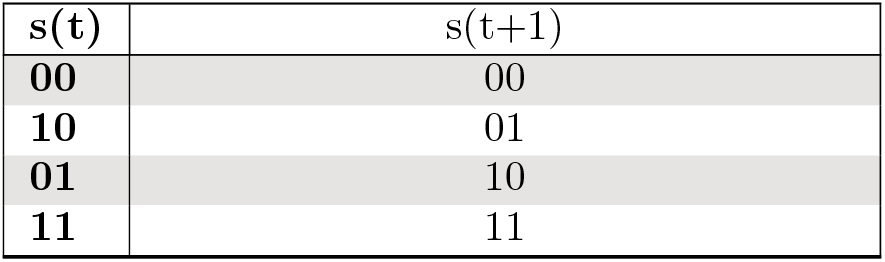
The transition probability matrix for a simple diode comprised of two interconnected `COPY` gates taking input from one another such as that described in Chalmers and McQueen [32]

## 6.4.2 AND+OR [23, 26]

Like the photodiode, the `AND+OR` system has been studied in the context of IIT at least twice prior to the current work [23, 26]. However, a concrete Φ value has yet to be published. Therefore, we take the “published value” to be that of the PyPhi value found in Section 2. Similarly, we take the initial state to be *s*_0_ = 00 in accordance with Section 2. The transition probability matrix is given below.

**Table 3:**
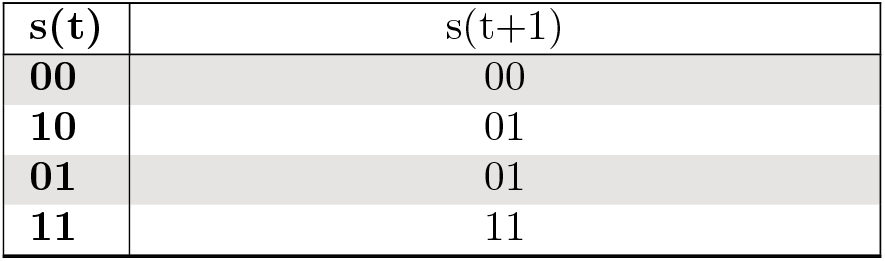
The transition probability matrix for an `AND+OR` system such as that described in Section 2

## 6.4.3 Hanson and Walker 2020 [23]

This system is a three bit digital counter in the initial state ’101’. The initial state is selected somewhat arbitrarily, since any initial state will work, but *s*_0_ = 101 results in a particularly fast evaluation. The TPM, from Figure 4 of the original publication, is as follows:

**Table 4:** Simple Electronic Counter Transition Probability Matrix from Hanson and Walker 2020 [24]

| $s(t)$ | $s(t+1)$ |
| --- | --- |
| 000 | 110 |
| 100 | 000 |
| 010 | 101 |
| 110 | 010 |
| 001 | 100 |
| 101 | 111 |
| 011 | 001 |
| 111 | 011 |

## 6.4.4 Majority Gate System

This system is comprised of three interconnected majority gates, each with three inputs, as shown in Figure 8. If the majority of inputs to a given node are 0 the state of the node at the next timestep is 0 and if the majority of inputs to a given node are 1 the state of the node at the next timestep is 1. In the main text, the system is evaluated in initial state *s*_0_ = 000. The transition probability matrix is provided below.

**Table 5:** `MAJ+MAJ+MAJ` System from Figure 8

| $s(t)$ | $s(t+1)$ |
| --- | --- |
| 000 | 000 |
| 100 | 000 |
| 010 | 000 |
| 110 | 111 |
| 001 | 000 |
| 101 | 111 |
| 011 | 111 |
| 111 | 111 |

## 6.4.5 Gomez et al. 2021 [20]

This papers studies the p53–Mdm2 biological regulatory network. Typically, this network is multivalued, but there are two possible binarizations that make standard Φ calculations possible. Of these, we chose the Fauré and Kaji binarization as it is much faster to analyze than the Tonello binarization. Following the authors, we choose an initial state *s*_0_ = 0001 and use the following TPM. Note, the PyPhi value we compute for this TPM differs from that published by the authors due to their use of several non-standard configuration settings, such as Krohn and Ostwalds definition of Φ as a difference in integrated conceptual information rather than the IIT 3.0 definition.

**Table 6:** Fauré-Kaji Binarization of the Transition Probability Matrix for the p53-Mdm2 Biological Regulatory Network from Gomez et al. 2021 [20]

| $s(t)$ | $s(t+1)$ |
| --- | --- |
| 0000 | 1101 |
| 1000 | 1100 |
| 0100 | 1100 |
| 1100 | 1110 |
| 0010 | 1101 |
| 1010 | 1101 |
| 0110 | 1101 |
| 1110 | 1111 |
| 0001 | 0001 |
| 1001 | 0000 |
| 0101 | 0000 |
| 1101 | 0010 |
| 0011 | 0001 |
| 1011 | 0001 |
| 0111 | 0001 |
| 1111 | 0011 |

## 6.4.6 Farnsworth 2021 [18]

In this paper a virocell (virus infected cell) is introduced into a Boolean network model of host cell dynamics. There are two network models provided, the first consists of five nodes and is the “full system”, while the second consists of three nodes and is the “reduced system”. For both systems, we study the case where all the nodes are ‘ON’ (i.e. *s*_0_ = 11111 and *s*_0_ = 111, respectively). Following the Supplementary Material provided by Farnsworth, the transition probabilities matrices are given below. Note, in the full system, the second node is an `AND` gate (as shown in Figure 6 of their main paper [18]) rather than a `COPY` gate as shown in Figure 8 of their Supplementary Material.

**Table 7:** The Transition Probability Matrix for the Entire Boolean Network Model of Virus-Host Dynamics from Farnsworth 2021 [18]

| s(t) | s(t+1) |
| --- | --- |
| 00000 | 00000 |
| 10000 | 00000 |
| 01000 | 10000 |
| 11000 | 10000 |
| 00100 | 01000 |
| 10100 | 01000 |
| 01100 | 11000 |
| 11100 | 11000 |
| 00010 | 00100 |
| 10010 | 00101 |
| 01010 | 10100 |
| 11010 | 10101 |
| 00110 | 01100 |
| 10110 | 01101 |
| 01110 | 11100 |
| 11110 | 11101 |
| 00001 | 00000 |
| 10001 | 00000 |
| 01001 | 10010 |
| 11001 | 10010 |
| 00101 | 01000 |
| 10101 | 01000 |
| 01101 | 11010 |
| 11101 | 11010 |
| 00011 | 00100 |
| 10011 | 00101 |
| 01011 | 10110 |
| 11011 | 10111 |
| 00111 | 01100 |
| 10111 | 01101 |
| 01111 | 11110 |
| 11111 | 11111 |

**Table 8:** The Transition Probability Matrix for the Reduced System from Farnsworth 2021 [18]

| s(t) | s(t+1) |
| --- | --- |
| 000 | 000 |
| 100 | 000 |
| 010 | 100 |
| Continued on next page |  |

| $s(t)$ | $s(t+1)$ |
| --- | --- |
| **110** | 101 |
| **001** | 010 |
| **101** | 010 |
| **011** | 110 |
| **111** | 111 |

## 6.4.7 Oizumi et al. 2014 [11]

This is the canonical `OR+AND+XOR` system that is often used in demonstrating how to calculate Φ [11, 15, 48]. Following Oizumi et al., we take the system to be in the initial state *s*_0_ = 100. The transition probability matrix is given below.

**Table 9:** The Transition Probability Matrix for the `OR+AND+XOR` system from Oizumi et al. 2014 [11]

| $s(t)$ | $s(t+1)$ |
| --- | --- |
| **000** | 000 |
| **100** | 001 |
| **010** | 101 |
| **110** | 100 |
| **001** | 100 |
| **101** | 111 |
| **011** | 101 |
| **111** | 110 |

## 6.4.8 Tononi et al. 2016 [22]

This paper demonstrates the calculation of Φ for a simple system of four interacting logic gates: `MAJORITY+OR+AND+AND`. Following the authors, we use the initial state *s*_0_ = 1110. The transition probability matrix is given below.

**Table 10:**
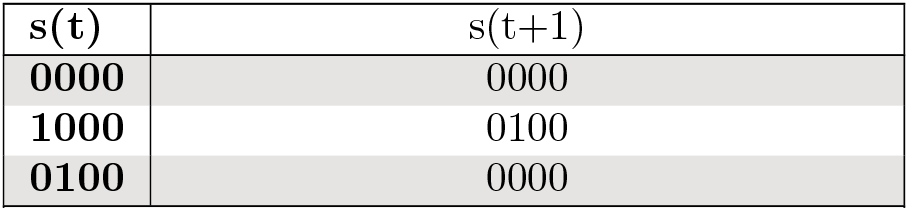
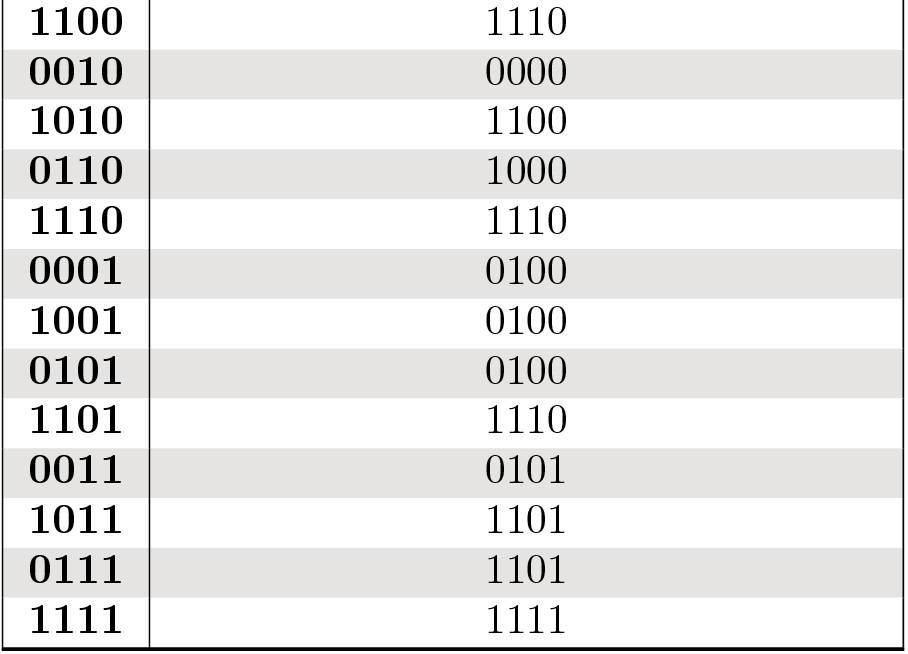
The Transition Probability Matrix for the `MAJ+OR+AND+AND` system from Tononi et al. 2016 [22]

| $s(t)$ | $s(t+1)$ |
| --- | --- |
| **0000** | 0000 |
| **1000** | 0100 |
| **0100** | 0000 |
| Continued on next page |  |
| 1100 | 1110 |
| 0010 | 0000 |
| 1010 | 1100 |
| 0110 | 1000 |
| 1110 | 1110 |
| 0001 | 0100 |
| 1001 | 0100 |
| 0101 | 0100 |
| 1101 | 1110 |
| 0011 | 0101 |
| 1011 | 1101 |
| 0111 | 1101 |
| 1111 | 1111 |

## 6.4.9 Hoel et al. 2016 [21]

This paper examines several small Boolean networks at both micro and macro scales. We choose to analyze the smallest of microsystems here, which is a system of four interconnected `AND` gates with noisy input. Following the authors, we analyze the system in initial state *s*_0_ = 0000. Due to the noisy input, the TPM is not deterministic and therefore cannot be written as an *N* by 2 matrix. Instead, it must be written as an *N* by *N* matrix where entry (*i, j*) specifies the probability of state *i* transitioning to state *j* at timestep *t* + 1 (a standard transition probability matrix). The transition probability matrix is given below.

**Table 11:** The Transition Probability Matrix for the noisy AND+AND+AND+AND system from Hoel et al. 2016 [21]

|  | 0 | 1 | 2 | 3 | 4 | 5 | 6 | 7 | 8 | 9 | 10 | 11 | 12 | 13 | 14 | 15 |
| --- | --- | --- | --- | --- | --- | --- | --- | --- | --- | --- | --- | --- | --- | --- | --- | --- |
| 0 | 0.24 | 0.10 | 0.10 | 0.04 | 0.10 | 0.04 | 0.04 | 0.02 | 0.10 | 0.04 | 0.04 | 0.02 | 0.04 | 0.02 | 0.02 | 0.01 |
| 1 | 0.24 | 0.10 | 0.10 | 0.04 | 0.10 | 0.04 | 0.04 | 0.02 | 0.10 | 0.04 | 0.04 | 0.02 | 0.04 | 0.02 | 0.02 | 0.01 |
| 2 | 0.24 | 0.10 | 0.10 | 0.04 | 0.10 | 0.04 | 0.04 | 0.02 | 0.10 | 0.04 | 0.04 | 0.02 | 0.04 | 0.02 | 0.02 | 0.01 |
| 3 | 0.00 | 0.00 | 0.00 | 0.00 | 0.00 | 0.00 | 0.00 | 0.00 | 0.00 | 0.00 | 0.00 | 0.00 | 0.49 | 0.21 | 0.21 | 0.09 |
| 4 | 0.24 | 0.10 | 0.10 | 0.04 | 0.10 | 0.04 | 0.04 | 0.02 | 0.10 | 0.04 | 0.04 | 0.02 | 0.04 | 0.02 | 0.02 | 0.01 |
| 5 | 0.24 | 0.10 | 0.10 | 0.04 | 0.10 | 0.04 | 0.04 | 0.02 | 0.10 | 0.04 | 0.04 | 0.02 | 0.04 | 0.02 | 0.02 | 0.01 |
| 6 | 0.24 | 0.10 | 0.10 | 0.04 | 0.10 | 0.04 | 0.04 | 0.02 | 0.10 | 0.04 | 0.04 | 0.02 | 0.04 | 0.02 | 0.02 | 0.01 |
| 7 | 0.00 | 0.00 | 0.00 | 0.00 | 0.00 | 0.00 | 0.00 | 0.00 | 0.00 | 0.00 | 0.00 | 0.00 | 0.49 | 0.21 | 0.21 | 0.09 |
| 8 | 0.24 | 0.10 | 0.10 | 0.04 | 0.10 | 0.04 | 0.04 | 0.02 | 0.10 | 0.04 | 0.04 | 0.02 | 0.04 | 0.02 | 0.02 | 0.01 |
| 9 | 0.24 | 0.10 | 0.10 | 0.04 | 0.10 | 0.04 | 0.04 | 0.02 | 0.10 | 0.04 | 0.04 | 0.02 | 0.04 | 0.02 | 0.02 | 0.01 |
| 10 | 0.24 | 0.10 | 0.10 | 0.04 | 0.10 | 0.04 | 0.04 | 0.02 | 0.10 | 0.04 | 0.04 | 0.02 | 0.04 | 0.02 | 0.02 | 0.01 |
| 11 | 0.00 | 0.00 | 0.00 | 0.00 | 0.00 | 0.00 | 0.00 | 0.00 | 0.00 | 0.00 | 0.00 | 0.00 | 0.49 | 0.21 | 0.21 | 0.09 |
| 12 | 0.00 | 0.00 | 0.00 | 0.49 | 0.00 | 0.00 | 0.00 | 0.21 | 0.00 | 0.00 | 0.00 | 0.21 | 0.00 | 0.00 | 0.00 | 0.09 |
| 13 | 0.00 | 0.00 | 0.00 | 0.49 | 0.00 | 0.00 | 0.00 | 0.21 | 0.00 | 0.00 | 0.00 | 0.21 | 0.00 | 0.00 | 0.00 | 0.09 |
| 14 | 0.00 | 0.00 | 0.00 | 0.49 | 0.00 | 0.00 | 0.00 | 0.21 | 0.00 | 0.00 | 0.00 | 0.21 | 0.00 | 0.00 | 0.00 | 0.09 |
| 15 | 0.00 | 0.00 | 0.00 | 0.00 | 0.00 | 0.00 | 0.00 | 0.00 | 0.00 | 0.00 | 0.00 | 0.00 | 0.00 | 0.00 | 0.00 | 1.00 |

## 6.4.10 Marshall et al. 2017 [16]

This model system is the fission yeast cell cycle from Marshall et al. 2017 [16]. As mentioned in the main text, we study the three node subsystem rather than the full eight node subsystem (plus one external node) studied in the original publication. To calculate the spectrum of Φ values for this subsystem (or just the PyPhi Φ value), the TPM for the entire system (all nine nodes) is required. Therefore, there are 512 states in the TPM. Following the authors, the initial state of the system is set to *s*_0_ = 000110011. In little-end binary notation (most significant bit on the right), the TPM used is as follows:

**Table 12:** Three Node Fission Yeast Transition Probability Matrix from Marshall et al. 2017 [16]

| **s(t)** | **s(t+1)** | **s(t)** | **s(t+1)** | **s(t)** | **s(t+1)** | **s(t)** | **s(t+1)** |
| --- | --- | --- | --- | --- | --- | --- | --- |
| **0** | 2 | **128** | 162 | **256** | 78 | **384** | 110 |
| **1** | 2 | **129** | 162 | **257** | 66 | **385** | 98 |
| **2** | 130 | **130** | 162 | **258** | 2 | **386** | 162 |
| **3** | 130 | **131** | 162 | **259** | 2 | **387** | 162 |
| **4** | 4 | **132** | 132 | **260** | 76 | **388** | 76 |
| **5** | 0 | **133** | 128 | **261** | 68 | **389** | 68 |
| **6** | 128 | **134** | 128 | **262** | 4 | **390** | 132 |
| **7** | 128 | **135** | 128 | **263** | 0 | **391** | 128 |
| **8** | 8 | **136** | 136 | **264** | 76 | **392** | 76 |
| **9** | 0 | **137** | 128 | **265** | 72 | **393** | 72 |
| **10** | 128 | **138** | 128 | **266** | 8 | **394** | 136 |
| **11** | 128 | **139** | 128 | **267** | 0 | **395** | 128 |
| **12** | 12 | **140** | 140 | **268** | 76 | **396** | 76 |
| **13** | 0 | **141** | 128 | **269** | 76 | **397** | 76 |
| **14** | 128 | **142** | 128 | **270** | 12 | **398** | 140 |
| **15** | 128 | **143** | 128 | **271** | 0 | **399** | 128 |
| **16** | 256 | **144** | 384 | **272** | 332 | **400** | 332 |
| **17** | 256 | **145** | 384 | **273** | 320 | **401** | 320 |
| **18** | 384 | **146** | 384 | **274** | 256 | **402** | 384 |
| **19** | 384 | **147** | 384 | **275** | 256 | **403** | 384 |
| **20** | 260 | **148** | 388 | **276** | 332 | **404** | 332 |
| **21** | 256 | **149** | 384 | **277** | 324 | **405** | 324 |
| **22** | 384 | **150** | 384 | **278** | 260 | **406** | 388 |
| **23** | 384 | **151** | 384 | **279** | 256 | **407** | 384 |
| **24** | 264 | **152** | 392 | **280** | 332 | **408** | 332 |
| **25** | 256 | **153** | 384 | **281** | 328 | **409** | 328 |
| **26** | 384 | **154** | 384 | **282** | 264 | **410** | 392 |
| **27** | 384 | **155** | 384 | **283** | 256 | **411** | 384 |
| **28** | 268 | **156** | 396 | **284** | 332 | **412** | 332 |
| **29** | 256 | **157** | 384 | **285** | 332 | **413** | 332 |
| **30** | 384 | **158** | 384 | **286** | 268 | **414** | 396 |
| **31** | 384 | **159** | 384 | **287** | 256 | **415** | 384 |
| **32** | 18 | **160** | 178 | **288** | 82 | **416** | 114 |
| **33** | 18 | **161** | 178 | **289** | 82 | **417** | 114 |
| **34** | 146 | **162** | 178 | **290** | 18 | **418** | 178 |
| **35** | 146 | **163** | 178 | **291** | 18 | **419** | 178 |
| Continued on next page |  |  |  |  |  |  |  |

| $s(t)$ | $s(t+1)$ | $s(t)$ | $s(t+1)$ | $s(t)$ | $s(t+1)$ | $s(t)$ | $s(t+1)$ |
| --- | --- | --- | --- | --- | --- | --- | --- |
| **36** | 16 | **164** | 144 | **292** | 84 | **420** | 84 |
| **37** | 16 | **165** | 144 | **293** | 80 | **421** | 80 |
| **38** | 144 | **166** | 144 | **294** | 16 | **422** | 144 |
| **39** | 144 | **167** | 144 | **295** | 16 | **423** | 144 |
| **40** | 16 | **168** | 144 | **296** | 88 | **424** | 88 |
| **41** | 16 | **169** | 144 | **297** | 80 | **425** | 80 |
| **42** | 144 | **170** | 144 | **298** | 16 | **426** | 144 |
| **43** | 144 | **171** | 144 | **299** | 16 | **427** | 144 |
| **44** | 16 | **172** | 144 | **300** | 92 | **428** | 92 |
| **45** | 16 | **173** | 144 | **301** | 80 | **429** | 80 |
| **46** | 144 | **174** | 144 | **302** | 16 | **430** | 144 |
| **47** | 144 | **175** | 144 | **303** | 16 | **431** | 144 |
| **48** | 272 | **176** | 400 | **304** | 336 | **432** | 336 |
| **49** | 272 | **177** | 400 | **305** | 336 | **433** | 336 |
| **50** | 400 | **178** | 400 | **306** | 272 | **434** | 400 |
| **51** | 400 | **179** | 400 | **307** | 272 | **435** | 400 |
| **52** | 272 | **180** | 400 | **308** | 340 | **436** | 340 |
| **53** | 272 | **181** | 400 | **309** | 336 | **437** | 336 |
| **54** | 400 | **182** | 400 | **310** | 272 | **438** | 400 |
| **55** | 400 | **183** | 400 | **311** | 272 | **439** | 400 |
| **56** | 272 | **184** | 400 | **312** | 344 | **440** | 344 |
| **57** | 272 | **185** | 400 | **313** | 336 | **441** | 336 |
| **58** | 400 | **186** | 400 | **314** | 272 | **442** | 400 |
| **59** | 400 | **187** | 400 | **315** | 272 | **443** | 400 |
| **60** | 272 | **188** | 400 | **316** | 348 | **444** | 348 |
| **61** | 272 | **189** | 400 | **317** | 336 | **445** | 336 |
| **62** | 400 | **190** | 400 | **318** | 272 | **446** | 400 |
| **63** | 400 | **191** | 400 | **319** | 272 | **447** | 400 |
| **64** | 66 | **192** | 194 | **320** | 78 | **448** | 78 |
| **65** | 66 | **193** | 194 | **321** | 66 | **449** | 66 |
| **66** | 130 | **194** | 130 | **322** | 66 | **450** | 194 |
| **67** | 130 | **195** | 130 | **323** | 66 | **451** | 194 |
| **68** | 68 | **196** | 196 | **324** | 76 | **452** | 76 |
| **69** | 64 | **197** | 192 | **325** | 68 | **453** | 68 |
| **70** | 128 | **198** | 128 | **326** | 68 | **454** | 196 |
| **71** | 128 | **199** | 128 | **327** | 64 | **455** | 192 |
| **72** | 72 | **200** | 200 | **328** | 76 | **456** | 76 |
| **73** | 64 | **201** | 192 | **329** | 72 | **457** | 72 |
| **74** | 128 | **202** | 128 | **330** | 72 | **458** | 200 |
| **75** | 128 | **203** | 128 | **331** | 64 | **459** | 192 |
| **76** | 76 | **204** | 204 | **332** | 76 | **460** | 76 |
| **77** | 64 | **205** | 192 | **333** | 76 | **461** | 76 |
| **78** | 128 | **206** | 128 | **334** | 76 | **462** | 204 |
| Continued on next page |  |  |  |  |  |  |  |

| $s(t)$ | $s(t+1)$ | $s(t)$ | $s(t+1)$ | $s(t)$ | $s(t+1)$ | $s(t)$ | $s(t+1)$ |
| --- | --- | --- | --- | --- | --- | --- | --- |
| 79 | 128 | 207 | 128 | 335 | 64 | 463 | 192 |
| 80 | 320 | 208 | 448 | 336 | 332 | 464 | 332 |
| 81 | 320 | 209 | 448 | 337 | 320 | 465 | 320 |
| 82 | 384 | 210 | 384 | 338 | 320 | 466 | 448 |
| 83 | 384 | 211 | 384 | 339 | 320 | 467 | 448 |
| 84 | 324 | 212 | 452 | 340 | 332 | 468 | 332 |
| 85 | 320 | 213 | 448 | 341 | 324 | 469 | 324 |
| 86 | 384 | 214 | 384 | 342 | 324 | 470 | 452 |
| 87 | 384 | 215 | 384 | 343 | 320 | 471 | 448 |
| 88 | 328 | 216 | 456 | 344 | 332 | 472 | 332 |
| 89 | 320 | 217 | 448 | 345 | 328 | 473 | 328 |
| 90 | 384 | 218 | 384 | 346 | 328 | 474 | 456 |
| 91 | 384 | 219 | 384 | 347 | 320 | 475 | 448 |
| 92 | 332 | 220 | 460 | 348 | 332 | 476 | 332 |
| 93 | 320 | 221 | 448 | 349 | 332 | 477 | 332 |
| 94 | 384 | 222 | 384 | 350 | 332 | 478 | 460 |
| 95 | 384 | 223 | 384 | 351 | 320 | 479 | 448 |
| 96 | 82 | 224 | 210 | 352 | 82 | 480 | 82 |
| 97 | 82 | 225 | 210 | 353 | 82 | 481 | 82 |
| 98 | 146 | 226 | 146 | 354 | 82 | 482 | 210 |
| 99 | 146 | 227 | 146 | 355 | 82 | 483 | 210 |
| 100 | 80 | 228 | 208 | 356 | 84 | 484 | 84 |
| 101 | 80 | 229 | 208 | 357 | 80 | 485 | 80 |
| 102 | 144 | 230 | 144 | 358 | 80 | 486 | 208 |
| 103 | 144 | 231 | 144 | 359 | 80 | 487 | 208 |
| 104 | 80 | 232 | 208 | 360 | 88 | 488 | 88 |
| 105 | 80 | 233 | 208 | 361 | 80 | 489 | 80 |
| 106 | 144 | 234 | 144 | 362 | 80 | 490 | 208 |
| 107 | 144 | 235 | 144 | 363 | 80 | 491 | 208 |
| 108 | 80 | 236 | 208 | 364 | 92 | 492 | 92 |
| 109 | 80 | 237 | 208 | 365 | 80 | 493 | 80 |
| 110 | 144 | 238 | 144 | 366 | 80 | 494 | 208 |
| 111 | 144 | 239 | 144 | 367 | 80 | 495 | 208 |
| 112 | 336 | 240 | 464 | 368 | 336 | 496 | 336 |
| 113 | 336 | 241 | 464 | 369 | 336 | 497 | 336 |
| 114 | 400 | 242 | 400 | 370 | 336 | 498 | 464 |
| 115 | 400 | 243 | 400 | 371 | 336 | 499 | 464 |
| 116 | 336 | 244 | 464 | 372 | 340 | 500 | 340 |
| 117 | 336 | 245 | 464 | 373 | 336 | 501 | 336 |
| 118 | 400 | 246 | 400 | 374 | 336 | 502 | 464 |
| 119 | 400 | 247 | 400 | 375 | 336 | 503 | 464 |
| 120 | 336 | 248 | 464 | 376 | 344 | 504 | 344 |
| 121 | 336 | 249 | 464 | 377 | 336 | 505 | 336 |
| Continued on next page |  |  |  |  |  |  |  |

| $s(t)$ | $s(t+1)$ | $s(t)$ | $s(t+1)$ | $s(t)$ | $s(t+1)$ | $s(t)$ | $s(t+1)$ |
| --- | --- | --- | --- | --- | --- | --- | --- |
| **122** | 400 | **250** | 400 | **378** | 336 | **506** | 464 |
| **123** | 400 | **251** | 400 | **379** | 336 | **507** | 464 |
| **124** | 336 | **252** | 464 | **380** | 348 | **508** | 348 |
| **125** | 336 | **253** | 464 | **381** | 336 | **509** | 336 |
| **126** | 400 | **254** | 400 | **382** | 336 | **510** | 464 |
| **127** | 400 | **255** | 400 | **383** | 336 | **511** | 464 |

## Footnotes

1 see *e.g.*, citation tracking statistics for IIT and other theories using the SCOPUS database [7]

2 Less than a day on a Mac Pro; 3.5 GHz 6-Core Intel Xeon E5 Processor

3 Implementation of the “Smallest”,“Biggest”, and “Moon” solutions are available via keyword arguments in the `PyPhi-Spectrum` package (see Algorithm 3), while the KO17 solution is available by changing the `USE-SMALL-PHI-FOR-CES` option in the standard `PyPhi` configuration file.

